# Kinesin family motors modify transcription mediated by ERR1 using a conserved nuclear receptor box motif

**DOI:** 10.1101/2022.11.08.515201

**Authors:** A.M.P.B Seneviratne, S. Lidagoster, S. Valbuena-Castor, K. Lashley, S. Saha, A. Alimova, Geri Kreitzer

**Author notes:** To whom correspondence should be addressed, Pramodh Seneviratne, PhD, Geri Kreitzer, PhD.

## Abstract

Kinesin family motors are microtubule (MT)-stimulated ATPases known best as transporters of cellular cargoes through the cytoplasm, regulators of MT dynamics, organizers of the mitotic spindle, and for insuring equal division of DNA during mitosis. Several kinesins have also been shown to regulate transcription by interacting with transcriptional cofactors and regulators, nuclear receptors, or with specific promotor elements on DNA. We previously showed that an LxxLL nuclear receptor box motif in the kinesin-2 family motor KIF17 mediates binding to the orphan nuclear receptor estrogen related receptor alpha (ERR1) and is responsible for the suppression of ERR1-dependent transcription by KIF17. Analysis of all kinesin family proteins revealed that multiple kinesins contain this LxxLL motif, raising the question as to whether additional kinesins motors contribute to regulation of ERR1. In this study, we interrogated the effects of multiple kinesins with LxxLL motifs on ERR1-mediated transcription. We demonstrate that the kinesin-3 motor KIF1B contains two LxxLL motifs, one of which binds to ERR1. In addition, we show that expression of a KIF1B fragment containing this LxxLL motif inhibits ERR1-dependent transcription by regulating nuclear entry of ERR1. We also provide evidence that the effects of expressing the KIF1B-LxxLL fragment on ERR1 activity are mediated by a mechanism distinct from that of KIF17. Because LxxLL domains are found in many kinesins, our data suggest an expanded role for kinesins in nuclear receptor mediated transcriptional regulation.

## Introduction

Kinesin family motors are microtubule stimulated ATPases that transport a large variety of cellular cargo along microtubules, regulate microtubule dynamics, and insure the equal segregation of genetic material during cell division (Hirokawa, Noda et al. 2009). In the human genome, there are 45 distinct kinesin genes. Apart from their canonical functions, a growing body of evidence highlights a non-canonical role for kinesins in regulation of transcription. Members of the kinesin-2 (KIF17) and kinesin-4 (KIF4, KIF7, Costal-2, BC12) families have well-documented roles, in diverse species and across phyla, in modifying the functions of transcription regulators or binding directly to specific promotor elements in DNA.

Mechanistically, the participatory role of kinesins in regulation transcription varies greatly. The kinesin KIF7 (Costal-2 in flies) sequesters Gli proteins (cubitis interruptus in flies) in the tip compartment of primary cilia (or cytoplasm in flies) and regulates their post-translational processing and transcriptional function in hedgehog signaling (Robbins, Nybakken et al. 1997, Sisson, Ho et al. 1997, Cheung, Zhang et al. 2009, Endoh-Yamagami, Evangelista et al. 2009, Liem, He et al. 2009, Marks and Kalderon 2011, He, Subramanian et al. 2014). The interaction of KIF7 with Gli proteins is mediated by a coiled-coil domain in KIF7 that mimics DNA and binds to zinc fingers in Gli (Haque, Freniere et al. 2022). KIF4 sequesters the DNA repair protein and transcriptional regulator poly-ADP-ribosyltransferase (PARP-1) in the nucleoplasm and inhibits its enzymatic activity until calcium influx is stimulated by membrane depolarization, resulting in CamKII-dependent release of PARP-1 from KIF4 (Midorikawa, Takei et al. 2006). The interaction of KIF4 with PARP-1 is mediated by the globular kinesin tail domain. Overexpression of a dominant-negative KIF4 tail domain in mouse neurons resulted in increased cell death, consistent with a role for PARP-1 in regulating anti-apoptotic gene expression. Kin4A in *O. sativa japonica* (also known as BC2 or GDD1) contains a bZIP motif and binds directly to the kaurene oxidase promotor. Kin4A-induced activation of *ent-*kaurene oxidase expression is required for the biosynthesis of gibberellic acid and cell elongation. Disruption of Kin4A binding to the kaurene oxidase promotor results in plant dwarfism (Li, Jiang et al. 2011). The testis specific mouse KIF17b regulates the cytoplasmic-nuclear shuttling of the activator of CREM mediated transcription (ACT) and CREM activity (Macho, Brancorsini et al. 2002). The interaction between KIF17b and ACT is mediated by an ∼100 amino acid region within the KIF17b stalk domain (Kotaja, Macho et al. 2005). In a complex, ACT-KIF17b translocate from the cytoplasm to the nucleus by a cAMP and PKA dependent but microtubule-independent mechanism.

We previously showed that the human kinesin 2 family member KIF17 interacts with the orphan nuclear receptor estrogen related receptor alpha (ERR1) (Seneviratne, Turan et al. 2017). Binding between KIF17 and ERR1 was mapped to the C-terminal tail domain of KIF17 and the C-terminal half of ERR1, which contains the ligand binding and activation factor 2 (AF2) domains (Seneviratne, Turan et al. 2017). The interaction of KIF17 with ERR1 is reduced significantly by deletion of 12 amino acids at the beginning of the KIF17 tail domain that includes a LxxLL nuclear receptor (NR) box. The LxxLL motif is conserved in transcriptional cofactors that bind nuclear receptors at AF2 domains (Bevan and Parker 1999) and is a multifunctional binding sequence involved in transcriptional regulation (Plevin, Mills et al. 2005). Thus, KIF17 appears to engage ERR1 in a manner similar to other regulatory cofactors. Consistent with this idea, expression of the KIF17 NR box in breast cancer cells inhibited the expression of a subset of ERR1 gene targets, suggesting that KIF17 is a selective, competitive inhibitor of ERR1 cofactor binding in the nuclear compartment.

ERR1 controls expression of genes involved in adaptive energy metabolism, cell proliferation, cell migration, cell differentiation, cell survival and autophagy, among others (Alaynick 2008, Audet-Walsh and Giguere 2015, Kim, Yang et al. 2018, Tripathi, Yen et al. 2020, Kim, Lee et al. 2021). As an orphan nuclear receptor, ERR1 is thought to be constitutively active. However, expression of its many gene targets is controlled, at least in part, by regulated binding to different transcriptional cofactors (Gaillard, Dwyer et al. 2007) (Kamei, Ohizumi et al. 2003, Schreiber, Knutti et al. 2003, Huss, Torra et al. 2004, Giguere, Dufour et al. 2011, Lu, Wang et al. 2011). With the discovery of cholesterol as a ligand for ERR1 (Wei, Schwaid et al. 2016) (Casaburi, Chimento et al. 2018), ERR1 could potentially be reclassified as an adoptive orphan. Because ERR1 expression and activity are also correlated with poor prognosis in many cancers (Ranhotra 2022) (Deblois, St-Pierre et al. 2013, Ranhotra 2015, Ranhotra 2022), ERR1 and identification of its potential modulators, is the subject of active research.

To determine if the mechanism used by KIF17 to regulate ERR1-dependent transcription is unique, we identified additional kinesin family members containing LxxLL motifs and assessed the effect of manipulating these kinesins on ERR1-mediated transcription. Here we show that a LxxLL domain present in the kinesin-3 family motor KIF1B interacts with and regulates ERR1-mediated transcription in breast cancer cells. This regulation leads to global suppression of ERR1 transcriptional targets, distinguishing it from the selective effects of KIF17 on ERR1 gene targets, and provides additional insight into the role of kinesins and the LxxLL motif in transcriptional regulation.

## Materials and Methods

### Cell culture, transfection, and microinjection

MCF7, MDA-MB-231 and HEK293T cells (ATCC) were cultured in DMEM (4.5 g/liter glucose, pH 7.2) with 5% FBS (Atlanta Biologicals/R&D Systems #S11150, Lot #F18071) 20 mM Hepes, and antibiotic/antimycotic (Fisher Scientific #15240062). Cells were incubated in a humidified environment with 5% CO2 at 37°C. For most experiments, expression constructs were transfected into cells using the Amaxa™ 4D-Nucleofector (10μg plasmid cDNA; Lonza, Basel, Switzerland) or JetPrime (2μg plasmid cDNA; Polyplus Transfection, Illkirch, France). For live-cell imaging experiments, cell nuclei were pressure injected with plasmid cDNAs (5-20μg/ml, details below) prepared in HKCl (10 mM Hepes, 140 mM KCl, pH 7.4) using back-loaded, glass capillary needles (Flaming/Brown micropipette puller, Sutter Instrument, Novado, CA) and a micromanipulator (MMO-202ND, Narishige, Greenville, NY) mounted on a Nikon TS100 inverted microscope.

### Expression constructs

KIF1A-NR, KIF1B-NR1, KIF1B-NR2, KIF18-NR, DNAs were amplified by PCR from human Caco2 cells or MCF7 breast cancer cells and cloned into Gateway™ expression vectors (Invitrogen) according to the manufacturer instructions. The KIF17-Tail and ERR1 constructs were described previously (Acharya, Espenel et al. 2013, Seneviratne, Turan et al. 2017). Primers used to generate additional expression constructs are:

KIF1A: forward; 5’ - GAC GCC ACC GAG CCT GCCand reverse; 5’ - GGA GAT TTT AGC AGT TCC CGA CTG GCG.

KIF1B-NR1: forward; 5’ - CCT GTG GAC TGG ACA TTT GCCand reverse; 5’ - AGA CCA ATA GTG TGT TGC TCC.

KIF1B-NR2: forward; 5’ - GAT CTC TTC AGT GAC GGG and reverse; 5’ - CAA GGA CAG ACC AGA ACG.

KIF18: forward; 5’ - GCT TGT CTT CAG GAA CAG CAA CAC AGG and reverse; 5’ - GGA CCA GTT CAG CCT ATT CCT TGT TGC.

ERRE-Luc reporter was provided by JM Vanacker (University of Lyon, France). 3X-ERE-Luc was purchased from Addgene (Plasmid 11354). KIF18A plasmids (Weaver, Ems-McClung et al. 2011) used to amplify selected regions of the protein were provided by Claire Walczak (Department of Biology, Indiana University).

All constructs were verified by sequencing prior to use in experiments.

### Co-immunoprecipitation and immunoblotting

HEK293T cells were transfected with indicated constructs in preparation for coimmunoprecipitation experiments. One day post transfection, cells were lysed in lysis buffer (50mM HEPES, pH 7.4 containing, 150mM NaCl, 1.5mM MgCl2, 0.5mM CaCl2, 10% (v/v) glycerol, 1% (v/v) Triton-X100, 1mM PMSF, and 0.5mg/ml each of leupeptin, bestatin, pepstatin) with rocking for 30 minutes. Lysates were incubated overnight at 4°C with 4-6μg rabbit anti-GFP antibody (Novus Biologicals, NB 600-303) or mouse anti-myc antibody (Sigma M4439) followed by immunoprecipitation with Protein-G sepharose beads (GE Healthcare). Other primary antibodies used for immunoblotting include mouse anti-GFP (Roche, 1814460) and rabbit anti-myc (Cell Signaling, 2276S).

### Luciferase reporter assays

Cells were transfected with indicated constructs. 24h after transfection cells were washed with PBS and lysed using Glo-lysis buffer (Promega) for 5min. Lysates were collected and mixed 1:1 with BrightGlo Luciferase assay reagent (Promega) and luminescence was measured using a luminometer (Molecular Devices, SpectraMax i3x). Results were corrected for background luminescence and plotted using GraphPad Prism 5. Statistical significance, *p* < 0.05, was determined using Bonferroni analysis.

### Time-lapse imaging and analysis

For live imaging of ERR1, cells were seeded onto heat-sterilized 25mm round coverslips 1-2 days prior to cDNA microinjection. Cells were co-injected with plasmids encoding ERR1-GFP (20ug/ml) and either mCh-KIF-NR (40ug/ml) or mCh-empty vector control (40ug/ml). After injections, cells were maintained in the incubator for 90min to allow for protein synthesis. Coverslips were then transferred to recording media (Hanks balanced salt solution containing 1% FBS, 4.5 g/L glucose, essential and nonessential amino acids, 20mM HEPES) and placed in a Sykes-Moore chamber (Bellco Glass, Vineland, NJ). The chamber was mounted in a temperature-controlled microincubator (PDMI-2, Harvard Apparatus) on a TE2000 inverted microscope running Elements™ acquisition and analysis package (Nikon Inc, Mellville, NY) and equipped with a Neo 5.5 cMOS camera (6.5um pixel, Andor Technologies). For MT depolymerization experiments, cells were treated with 33μM nocodazole (NZ; Sigma M1404) immediately following cDNA injection and maintained in NZ for the duration of the experiment. Time-lapse images were acquired at 10-minute intervals for 3 hours using a 20x (NA 0.75) plan apochromat objective. For image analysis, background fluorescence was subtracted and regions of interest (ROIs) were identified using morphometric thresholding around individual cells and their corresponding nuclei. Integrated florescence intensities were measured for nuclear ROIs (N) and total cell area ROIs (T), and expressed as a ratio, N/T fluorescence per cell over time. Average N/T fluorescence per cell at the beginning and end of the time-lapse were plotted using Graphpad Prism 5.

### qRT-PCR

qRT-PCR was performed using SyBr green on a QuantStudio 7 Flex Real-time qPCR system using standard methods. Total RNA and cDNA were obtained from stable cell lines using Qiagen RNAeasy RNA extraction kit and cDNA generated using standard methods. Tbp1 was used as a control. All primers were obtained from Azenta/Genewiz. The following primers were used:

Tbp1: forward; 5’ –TGT ATC CAC AGT GAA TCT TGG TTG and reverse; 5’ –GGT TCG TGG CTC TCT TAT CCT C

14-3-3: forward; 5’ – GCATGAAGTCTGTAACTGAGCA and reverse; 5’ – GCACCTTCCGTCTTTTGTTC

Claudin-4: forward; 5’ –CCATATAACTGCTCAACCTGTCC and reverse; 5’ – AGATAAAGCCAGTCCTGATGC

ERR1: forward; 5’ –TCTCCGCTTGGTGATCTCA and reverse; 5’ – CTATGGTGTGGCATCCTGTG

HIF1A: forward; 5’ –CAACCCAGACATATCCACCTC and reverse; 5’ – CTCTGATCATCTGACCAAAACTCA.

HIF2: forward; 5’ –CTTTGCGAGCATCCGGTA and reverse; 5’ – AGCCTATGAATTCTACCATGCG

Osteopontin: forward; 5’ –CCCCACAGTAGACACATATGATG and reverse; 5’ – TTCAACTCCTCGCTTTCCAT

PGC1A: Forward 5’ GTC CTT TTC TCG ACA CAG GT and reverse GTC TGT AGT GGC TTG ACT CAT AG.

## Results

### Identification of kinesin family motor transcriptional regulator candidates

To assess if kinesin family motors other than KIF17 have the potential to act as transcriptional regulators of ERR1 in the nucleus, we searched the sequences of all kinesin family motors in the human genome to identify those which contain LxxLL motifs. We then selected candidates for experimental assessment based on (i) the location of the LxxLL motif within the entire protein sequence, and (ii) the presence and predicted strength of nuclear localization signals (NLS). We eliminated kinesins that contained LxxLL motifs solely in the microtubule and ATP-binding motor domain. We also eliminated kinesins that lacked at least one predicted NLS outside the motor domain. The remaining sequences were then sorted by the predicted strength of the NLS using NLS mapper (https://nls-mapper.iab.keio.ac.jp/cgi-bin/NLS_Mapper_form.cgi), where only kinesins with NLS sequences predicted to result in strong nuclear localization, or localization to both the cytoplasm and nucleus (Kosugi, Hasebe et al. 2009, Kosugi, Hasebe et al. 2009) were considered further. Kinesins that did not contain classical monopartite or bipartite NLS sequences (Lu, Wu et al. 2021) were also eliminated. The kinesins, KIF17, KIF18A, KIF1A and KIF1B met both these criteria. KIF17, KIF18A, KIF1A contained 1 LxxLL motif while KIF1B contained 2 LxxLL motifs (denoted KIF1B-NR1 and KIF1B-NR2) (Figure 1).

**Figure 1.**
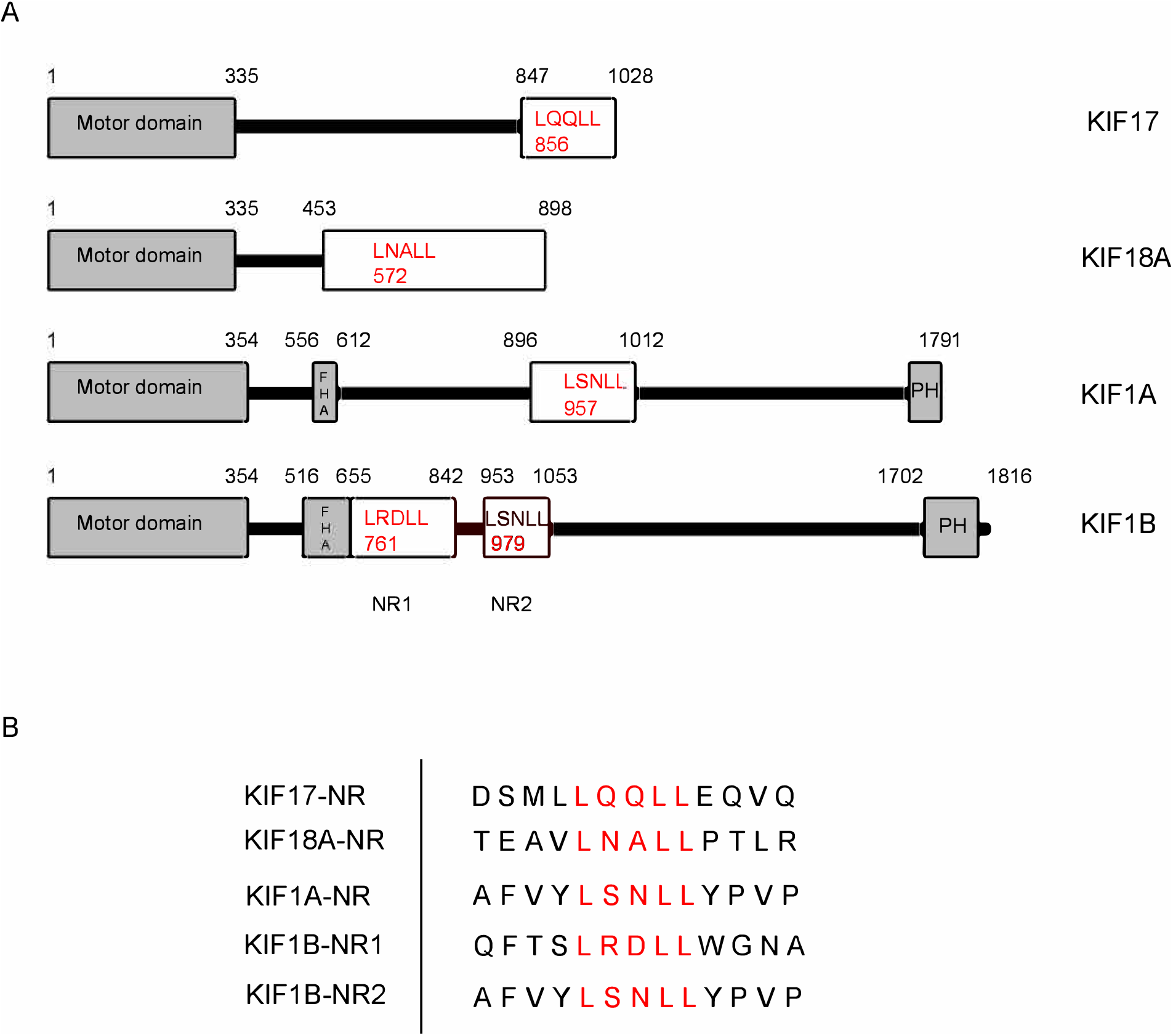
Kinesin family motors contain LxxLL nuclear receptor box motifs. (A) Schematic diagrams showing the kinesins with LxxLL motifs used in this study. The kinesin motor domains are indicated by the grey boxes. Kinesin fragments used to generate expression plasmids are shown in the white boxes. LxxLL motifs within these fragments are shown in red text, with numbers indicating the position of the first leucine of the motif. The numbers above each diagram represent the amino acid positions within the full-length kinesin sequence. (B) Sequence alignment of LxxLL domains of the kinesin constructs used in the study. Red lettering represents the LxxLL motifs.A

### KIF1B-NR1 inhibits of ERR1-dependent transcription

We designed epitope or fluorescently tagged expression plasmids encoding fragments of KIF1A, KIF1B, and KIF18A that contained the LxxLL motifs identified in in these kinesins (denoted *tag*-KIF*X*-NR). We next co-expressed these kinesin fragments in MCF7 breast cancer cells with a plasmid containing the ERR1 responsive element fused to luciferase (ERRE-Luc) and performed luminescence reporter assays to determine if expression of the LxxLL regions of these kinesins alters ERR1-dependent transcription. As a negative control, we expressed either myc- or GFP-empty vector (denoted *tag*-EV). As a positive control, we expressed the NR box-containing KIF17 tail (GFP-KIF17-T), which we showed to inhibit ERR1-mediated transcription in MCF7 and MDA-MB-231 cells (Seneviratne, Turan et al. 2017). Expression of GFP-KIF17-T reduced luciferase production by 35%, consistent with our prior study. Expression of GFP-KIF1B-NR1 reduced luciferase production by ERRE-Luc by 50% of empty vector control (Figure 2A). By contrast, expression of GFP-KIF1A-NR, KIF1B-NR2 and KIF18A-NR had no significant impact on luciferase production from the ERRE-Luc reporter. These data suggest that KIF1B, through its NR1-containing region, may act as a second regulator of ERR1-mediated transcription.

**Figure 2.**
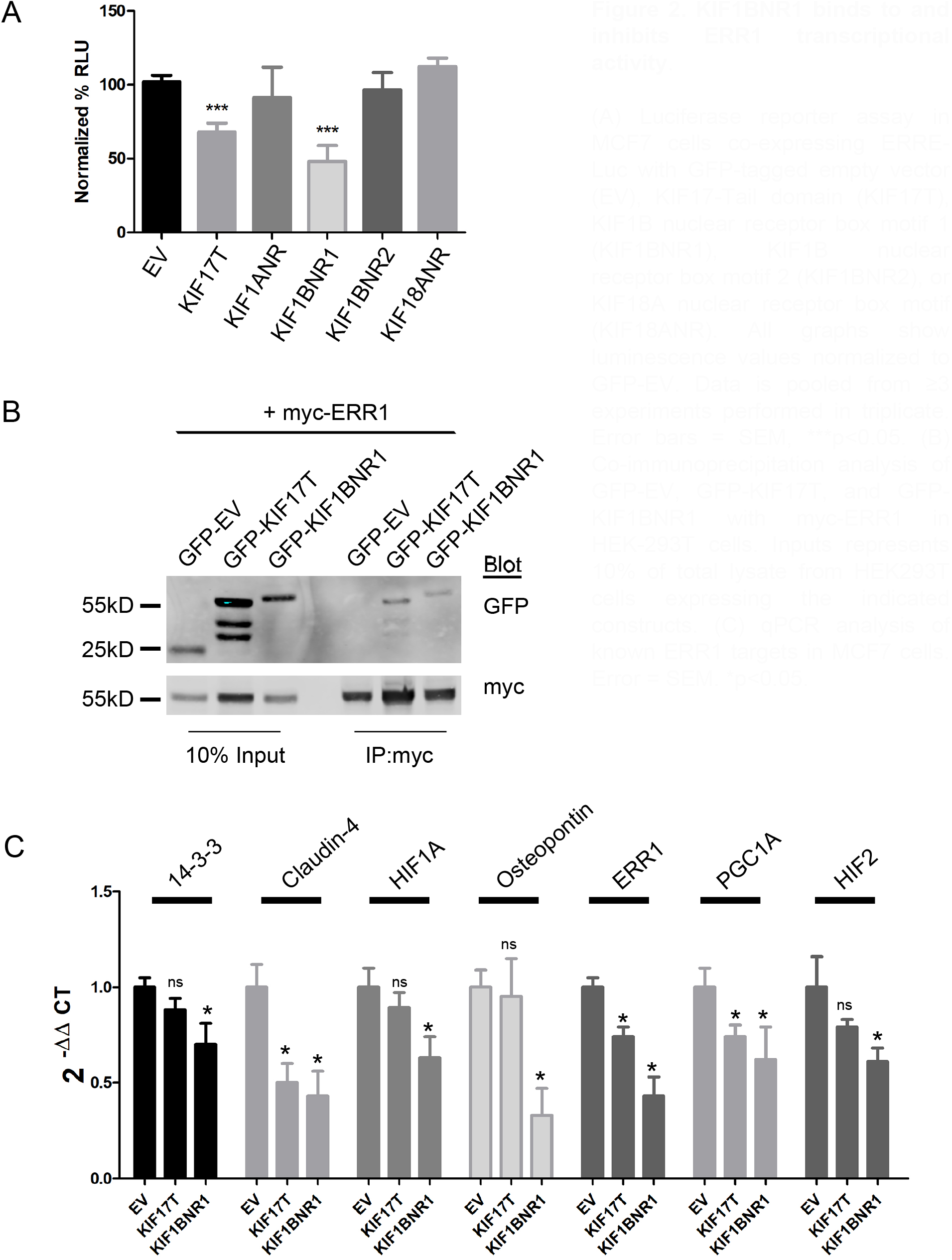
KIF1BNR1 binds to and inhibits ERR1 transcriptional activity. (A) Luciferase reporter assay in MCF7 cells co-expressing ERRE-Luc with GFP-tagged empty vector (EV), KIF17-Tail domain (KIF17T), KIF1B nuclear receptor box motif 1 (KIF1BNR1), KIF1B nuclear receptor box motif 2 (KIF1BNR2), or KIF18A nuclear receptor box motif (KIF18ANR). All graphs show luminescence values normalized to GFP-EV. Data is pooled from ≥3 experiments performed in triplicate. Error bars = SEM, ***p<0.05. (B) Co-immunoprecipitation analysis of GFP-EV, GFP-KIF17T, and GFP-KIF1BNR1 with myc-ERR1 in HEK-293T cells. Inputs represents 10% of total lysate from HEK293T cells expressing the indicated constructs. (C) qPCR analysis of known ERR1 targets in MCF7 cells. Error = SEM. *p<0.05.

### KIF1B-NR1 interacts with ERR1

We performed co-immunoprecipitation analysis to determine if the effects of KIF1B-NR1 on ERR1-dependent transcription results from an interaction of these proteins. We co-expressed myc-ERR1 and GFP-KIF1B-NR1 or GFP-KIF1B-NR2 in HEK293T cells. One day after transfection, we lysed the cells, reserved 10% of this total lysate, and incubated the remainder of cell lysates with anti-myc antibody to immunoprecipitate myc-ERR1. In these experiments, we found that GFP-KIF1B-NR1 coprecipitates with myc-ERR1, while GFP-KIF1B-NR2 does not (Figure 2B). This data shows that although KIF1B contains 2 LxxLL NR box motifs, only one of these, the NR1, can bind to ERR1. Thus, the effects of KIF1B-NR1 on ERR1-dependent transcription are likely mediated by binding of KIF1B-NR1 to ERR1.

### KIF1B-NR1 globally inhibits ERR1 target genes

We showed previously that expression of the KIF17 tail domain inhibits transcription of a subset of the known ERR1 regulated gene targets (Seneviratne, Turan et al. 2017). Since expression of either KIF1B-NR1 or KIF17-T inhibits ERR1-dependent transcription in reporter assays, we next tested if expression of KIF1B-NR1 inhibits the same or different gene targets of ERR1 using qPCR. For these experiments, we generated MCF7 stable cell lines expressing either GFP-EV as a negative control, GFP-KIF1B-NR1 or GFP-KIF17-T. We then compared the effects KIF1B-NR1 with that of KIF17-T and empty vector on the expression of 7 genes under the control of ERR1. Expression of KIF1B-NR1 significantly inhibited mRNA expression of all 7 gene targets tested (Figure 2C). By contrast, and consistent with our prior studies in MDA-MB-231 breast cancer cells (Seneviratne, Turan et al. 2017), expression of KIF17-T selectively inhibited expression of ERR1 targets, including ERR1, PGC1A and claudin-4, but had no significant effect on the mRNA levels of 14-3-3ζ, HIF1A, HIF2 and osteopontin. These data demonstrate that although both KIF1B-NR1 and KIF17-T bind ERR1 and inhibit its transcriptional activity in reporter assays, they do not likely act by the same mechanisms since their effects on transcription of specific ERR1-dependent gene targets are distinct.

### KIF1B-NR1 inhibits nuclear import of ERR1 by sequestering ERR1 in the cytoplasm

The non-selective inhibition of ERR1 target gene expression by KIF1B-NR1 suggests that it may act as a dominant-negative inhibitor that sequesters ERR1 from DNA. To test this idea, we used time-lapse fluorescence microscopy to monitor the subcellular distribution of newly synthesized ERR1 in the presence of empty vector control or KIF1B-NR. We co-expressed mCh-ERR1 and GFP-EV or GFP-KIF1B-NR1 in MCF-7 cells by intra-nuclear injection of plasmid DNA. 90 minutes after microinjection, the protein products of these cDNAs could be detected microscopically. At this time, cells were transferred to a temperature-controlled chamber on the microscope and the distribution of mCh-ERR1 and GFP-EV or GFP-KIF1B-NR1 was monitored at 10-minute intervals for 3 hours at 35°C (Figure 3A). From these images, we quantified the cytoplasmic and nuclear ERR1 fluorescence over time to determine the impact of expressing KIF1B-NR1 on translocation of newly synthesized mCh-ERR1 from the cytoplasm to the nucleus (Figure 3B). At the beginning of all recordings we routinely measured ∼20-30% of the total mCh-ERR1 in the nucleus. In control cells expressing GFP-EV and mCh-ERR1, the fraction of nuclear mCh-ERR1 increased by ≥ 2-fold over the course of 3h. Conversely, in cells expressing mCh-ERR1 and GFP-KIF1B-NR, the translocation of mCh-ERR1 from the cytoplasm to the nucleus was inhibited nearly completely. As expected, expression of GFP-KIF1B-NR2, which does not bind ERR1, had no effect on nuclear translocation of co-expressed mCh-ERR1 in these assays.

**Figure 3.**
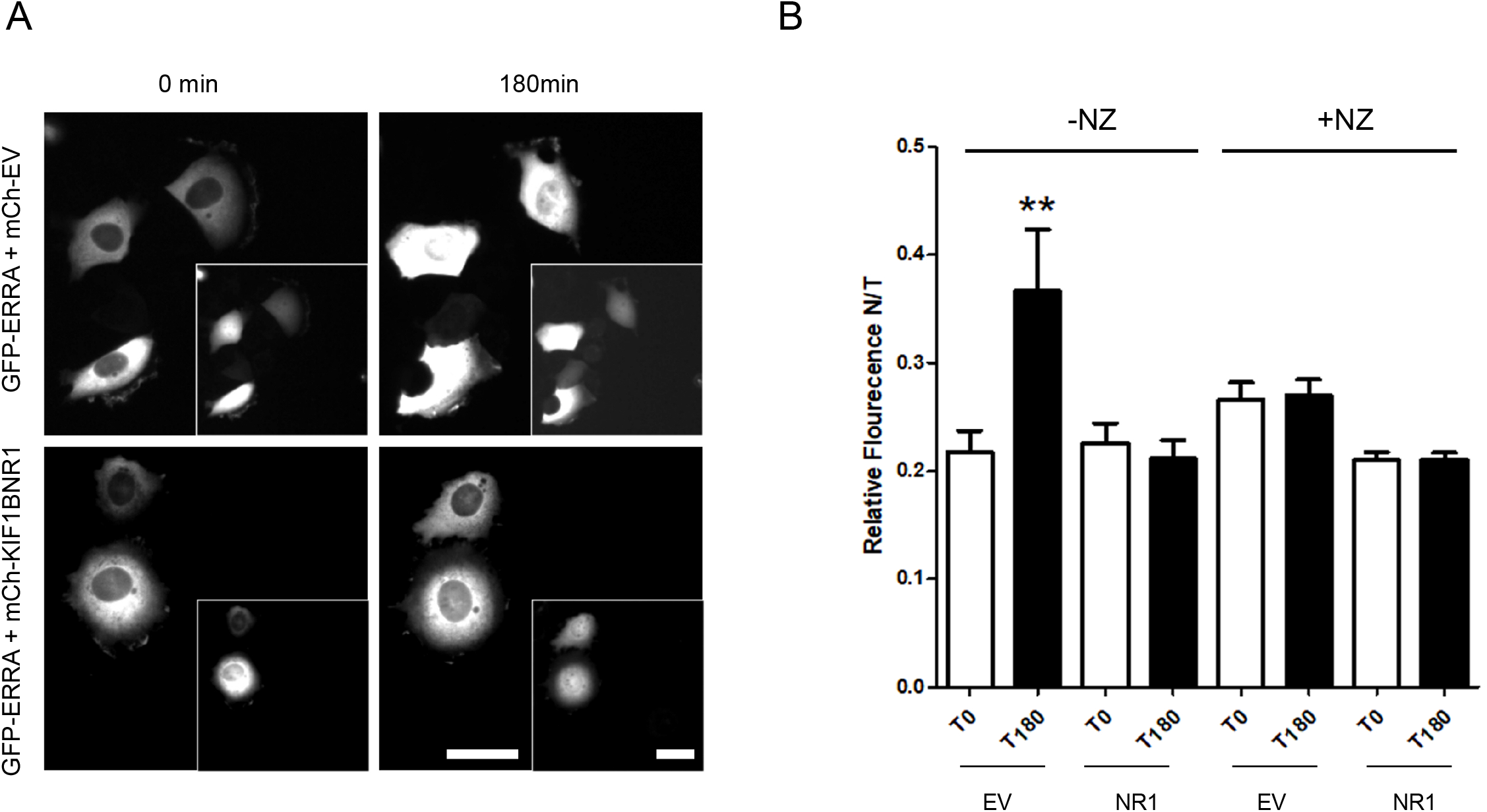
Nuclear accumulation of newly expressed ERR1 is inhibited by expression of KIF1BNR1 and is independent of an intact microtubule network. (A) MCF-7 cells injected with GFP-ERR1 and control, mCh-EV or mCh-KIF1BNR1 cDNAs. Images show GFP-ERR1 in first and last frames of time-lapse recordings. Insets: mCh-EV control and mCh-KIF17-T in injected cells. Scale bars = 20μm. (B) Graphs showing the fraction of nuclear GFP-ERR1, as a percentage of total cellular GFP-ERR1 (N/T fluorescence), in cells co-expressing mCh-EV or mCh-KIF1BNR1. GFP-ERR1 fluorescence intensities (nuclear and total) were quantified at the first and last time points in recordings of cells with intact microtubules (-NZ), and in cells where microtubules are disrupted (+NZ). Error bars = SEM. ** p<0.05.

To determine if the effects KIF1B-NR1 on nuclear import of ERR1 are dependent on an intact MT network, we treated cells with nocodazole to depolymerize MTs. Nocodazole was added to cells immediately after cDNA injection, and cells were continuously incubated in the drug during the 3h time-lapse imaging. Similar to untreated controls, expression of GFP-KIF1B-NR1 also inhibited nuclear import of newly synthesized mCh-ERR1 in the presence of nocodazole. This data indicates that KIF1B-NR1 sequesters ERR1 in the cytoplasm in a MT-independent manner. The MT-independent sequestration of ERR1 in the cytoplasm by KIF1B-NR1 contrasts the MT-dependent sequestration of ERR1 shown for cells expressing the KIF17 tail domain (Seneviratne, Turan et al. 2017).

## Discussion

Many kinesin family motors encode one or more LxxLL motifs, a sequence that is conserved in nuclear receptor coactivator proteins and is known as the nuclear receptor box motif. The function of this motif in kinesins, however, is not known. Further, although roles for several kinesins in regulation of transcription have now been documented, the function of kinesin LxxLL motifs in transcriptional regulation has gone largely unexplored. We showed previously that the LxxLL motif contained within the tail domain of KIF17 is both necessary and sufficient to modify expression of ERR1-dependent genes in breast cancer cells (Seneviratne, Turan et al. 2017). In our prior work, we also demonstrated that KIF17 regulates ERR1 function in a manner which suggests it competes with nuclear receptor transcriptional co-activators in the nucleus. The goal of this study was to assess if LxxLL motifs encoded by additional kinesin family motors are also able to engage ERR1 and modify its activity.

The presence of one or more LxxLL motifs in numerous kinesin family members made it necessary to prioritize the kinesin motors to be examined using stringent criteria. First, since the microtubule and ATP-binding domains of all kinesins are highly conserved (Hirokawa and Takemura 2004) and are required to execute the motor functions of these proteins, we hypothesized that only LxxLL domains outside the kinesin motor domain would be functionally relevant in engaging transcription regulators. LxxLL motifs located in known or predicted kinesin dimerization domains were also eliminated from consideration. Second, because nuclear receptor function and transcriptional regulation occurs in the nucleus, we limited our search to kinesins encoding a strong NLS outside the kinesin motor domain. Non-classical NLS (Lu, Wu et al. 2021) were also eliminated, but should be considered further in future studies. For example, KIF11, a kinesin involved in separation of spindle poles during mitosis (Kapitein, Peterman et al. 2005), was shown to interact with thyroid hormone receptor beta in yeast-2-hybrid assays, and with NCoR1 in affinity-MS proteomic assays (Fozzatti, Lu et al. 2011). Although the functional significance of these interactions are not yet known, KIF11 possesses an LxxLL motif outside of its motor domain, but it does not possess a classical NLS and, instead, encodes a non-classical, long linker (20 amino acid), bipartite NLS of moderate strength. As there is no evidence to date indicating KIF11 localizes to the nucleus in interphase cells, we choose to eliminate it as a candidate in our study. However, a broader consideration of kinesin LxxLL domains, along with rigorous examination of kinesin localization to the nucleus in interphase, could justify further investigation of KIF11 and other motors with respect to transcriptional regulation.

Kinesin motors that conformed best to our criteria included KIF17, KIF18A, KIF1A and KIF1B. Consistent with our prior work, expression of the LxxLL containing KIF17-Tail domain inhibited ERR1 dependent transcription. Of the other kinesins tested, ERR1 dependent transcription was affected only by expression of the KIF1B-NR1. Notably, KIF1B-NR2 had no impact on ERR1 function. The LxxLL motifs in KIF1B-NR1 (LRDLL) and KIF1B-NR2 (LSNLL) are both consistent with Class I nuclear receptor box motifs, which contain 1 polar amino acid residue immediately preceding the first leucine in the motif (Savkur and Burris 2004). This suggests that, rather than sequence differences, the differential actions of these LxxLL domains might be due to surface accessibility of the LxxLL motifs. Sequence analysis of KIF1B NR1 and NR2 shows that both these LxxLL motifs reside outside the FHA and PH domains of the protein. No structure for full length KIF1B, or for the relevant regions of KIF1B examined here currently exist. However, predictive structures of KIF1B NR1 and NR2 regions in AlphaFold (Jumper, Evans et al. 2021) (https://alphafold.ebi.ac.uk/entry/O60333), and analysis of solvent accessible surface area, indicate that the KIF1B-NR1 is more exposed than that of KIF1B-NR2. Thus, the conformation of these regions may contribute to the ability of these domains to exert effects on ERR1. The LxxLL motif and surrounding amino acids in KIF1A is identical to that of the KIF1B-NR2. As such, the lack of impact on ERR1 function was not surprising, and could also be due to inaccessibility. Alternatively, the amino acids located further upstream and downstream of the LxxLL motifs might be important in regulating binding to ERR1, as described for some transcriptional coactivators. This possibility will need to be tested in future work.

The kinesin-8 family motor KIF18A also contains LxxLL motifs, and was reported to bind estrogen receptors in the cytoplasm of stromal cells in the presence of estradiol (Zusev and Benayahu 2009). However, the interaction between KIF18A and estrogen receptor was not mapped, and thus the significance of the LxxLL motif to KIF18A and estrogen receptor binding is not known. Furthermore, the effect of KIF18A on estrogen receptor dependent transcription was not reported, so it is unclear if KIF18A acts to modify estrogen receptor-mediated gene expression. In our assays, expression of KIF18A fragments containing the LxxLL domain had no impact on ERR1-mediated transcription. As such, it is not yet clear whether KIF18A LxxLL motifs are functionally significant.

Our prior work, and the data shown here, suggest at least two possible mechanisms by which the interaction of a kinesin with ERR1 regulates ERR1 function. KIF1B regulates ERR1-dependent transcription globally by sequestering ERR1 in the cytoplasm. Although the effect of expressing KIF1B-NR on transit of ERR1 to the nucleus is MT-independent, we don’t yet know if regulation of ERR1 localization by native KIF1B (or expressed, full-length KIF1B) depends on MTs, or if KIF1B interacts with ERR1 in its monomeric or dimeric form (Nangaku, Sato-Yoshitake et al. 1994, Soppina, Norris et al. 2014). KIF17 regulates ERR1-dependent transcription selectively, by modifying ERR1 function in the nucleus. Interestingly, the subset of ERR1-dependent genes affected by expression of KIF17-Tail in MCF-7 differed from those affected in MDA-MB-231 cells (Seneviratne, Turan et al. 2017). The differential effects of KIF17 on expression of ERR1 gene targets could be due to the differential activation of gene expression by ERR1 as compared with those genes activated by ERR1-ER heterodimers in cells that also express estrogen receptor, as in MCF-7 (Yang, Shigeta et al. 1996, Vanacker, Pettersson et al. 1999, Kraus, Ariazi et al. 2002). MDA-MB-231 are triple-negative breast cancer cells, and thus do not express estrogen receptor. The selective effects of expressing KIF17-Tail on ERR1-dependent transcription could also be due to displacement of a subset of ERR1 transcriptional coactivators which may be differentially expressed in ER positive and ER-negative cells.

In summary, the work presented here, along with prior studies documenting a role for kinesin motors in regulating gene transcription, highlight gaps in our knowledge of non-canonical kinesin functions. However, it is clear that kinesins use multiple mechanisms to exert effects on the activity of transcription factors. Further investigation will be needed to determine if additional nuclear receptors are regulated by kinesins, and if LxxLL motifs in additional kinesin family motors are utilized to regulate gene expression more broadly. If so, LxxLL sequences could be envisioned as tools to manipulate expression of ERR1 gene targets pharmacologically and selectively.

## Acknowledgments

This work was supported by new lab start-up funds to GK from the CUNY School of Medicine and City College of New York. SL was supported in part by NSF-REU #1852496 and by a Jack Rudin and Lewis Rudin Research Fellowship.

